# Bi-directional relationship between the biofilm of *Porphyromonas gingivalis* and the amyloid-beta peptide

**DOI:** 10.1101/2025.05.16.654480

**Authors:** David Dumoulin, Mnar Ghrayeb, Sarah Côté, Daniel Garneau, Liraz Chai, Eric H. Frost, Tamàs Fülöp, Pascale B. Beauregard

## Abstract

Periodontitis and *Porphyromonas gingivalis* infections are significant risk factors for the onset of Alzheimer’s disease (AD). Despite *P. gingivalis* relying on biofilm for its survival and virulence, the impact of the extracellular matrix on AD’s neuropathological hallmarks was never examined. In this study, we report a bidirectional relationship between the amyloid beta (Aβ) peptide, which plays a central role in AD, and the biofilm of *P. gingivalis*. Using multiple fluorescent markers for biofilm components, we observed that Aβ1-40 inhibited biofilm formation while Aβ1-42 increased extracellular matrix production. Also, using thioflavin T staining and atomic force microscopy, we observed co-aggregation between the biofilm and monomeric Aβ1-40, resulting in a quicker aggregation and significant changes in aggregate structures. Our findings propose mechanistic explanations for the role of *P. gingivalis* as a risk factor for AD and offer potential mechanisms for the microbial involvement in AD etiology.

**Importance:** While the etiology of Alzheimer’s disease has been studied extensively for the past 50 years, its exact causes remain unknown. Our current understanding is that the accumulation of multiple genetic and environmental risk factors would lead to the onset of the disease. *Porphyromonas gingivalis* is a bacterium that produces biofilm and elicits periodontitis, a chronic infection of the gums that constitutes a risk factor for Alzheimer’s disease. While studies have looked at the effects of *P. gingivalis* in triggering Alzheimer’s symptoms in animal models, none have explored the impact of the biofilm, which is ubiquitous to this bacterium. Our study seeks to bridge that gap by demonstrating a bi-directional relationship between the biofilm of *P. gingivalis* and amyloid beta, one of the brain lesions involved in Alzheimer’s. By understanding risk factors involved in Alzheimer’s and their impact, we hope to provide valuable knowledge on prevention and treatment.

## Introduction

The oral biofilm is a complex microbial community involved in the onset of multiple diseases, including periodontitis. Periodontitis is a chronic and inflammatory condition of the periodontium characterized by a bacterial-driven degradation of the gum epithelium and the underlying alveolar bone tissue^1^. Onset of periodontitis is usually associated with proliferation within the gingival biofilm of bacteria from the red complex, *Treponema denticola, Tannerella forsythia* and *Porphyromonas gingivalis*^2,3^. *P. gingivalis* specifically contributes to the establishment of the oral biofilm by suppressing the host’s immune response and releasing nutrients through periodontal tissue breakdown^4,5^. *P. gingivalis* produces gingipains, cysteine proteases targeting lysine-Xaa (Kgp) or arginine-Xaa (RgpA, RgpB) bonds, that contribute to intracellular invasion and degradation of antimicrobial peptides^6–10^.

Being strictly anaerobic, *P. gingivalis* depends on its biofilm to limit O2 exposure. This lifestyle allows for its survival and the secretion of virulence factors. Although poorly characterized, *P. gingivalis* biofilm relies on the production of two co-occurring fimbriae, the major fimbria FimA and the minor fimbria Mfa1^11,12^, as well as on exopolysaccharides production^13^. *P. gingivalis* biofilm production also correlates with gingipain secretion, thus contributing to epithelial tissue breakdown, cellular invasion, and release in the bloodstream^14,15^. *P. gingivalis* propagation influences the course of multiple diseases such as atherosclerosis^16^, rheumatoid arthritis (reviewed in^17^), and adverse pregnancy outcomes (reviewed in^18^). Periodontitis and *P. gingivalis* infections are also confirmed risk factors for Alzheimer’s disease (AD)^19^.

AD is an incurable neurodegenerative disease characterized by the accumulation of aggregated amyloid-beta (Aβ) plaques and intracellular tangles of hyperphosphorylated Tau proteins. The occurrence of these lesions’ triggers neuroinflammation and neuronal cell death, leading to cognitive decline in older adults^20^. The toxicity of Aβ is linked to amyloid aggregation into oligomers, which form pores in neuronal cell membranes^21,22^, causing mitochondrial dysfunction^23^ and triggering pathogenic inflammation in the central nervous system^24^. Aβ is mainly secreted under two forms: Aβ1-40 represents ∼90% of the overall Aβ pool but exhibits low toxicity and aggregation; Aβ1-42 is highly aggregative, neurotoxic, and overrepresented in Aβ plaques^25^.

The involvement of microbial infection in AD etiology has been the subject of increasing interest in the last decade^26^. Associations between AD and diverse infectious agents revealed that Aβ acts as an antimicrobial peptide against viral, fungal and bacterial pathogens^27–31^. Indeed, in response to different infections, Aβ1-42 is strongly produced and accumulates in the central nervous system (CNS; review in^28^), hinting toward a mechanism in which infections would contribute to AD etiology. Of note, AD development occurs 20-30 years before the occurrence of symptoms^32^, which suggests the involvement of a chronic infection instead of an acute event.

*P. gingivalis* is one of the most studied pathogens concerning AD, since many of its biomolecules, including nucleic acids, lipopolysaccharides (LPS) and gingipains, were observed in various regions of AD brains (reviewed in^33^). *P. gingivalis* was also isolated from AD patients’ cerebrospinal fluid^34,35^. In mouse models, repeated oral infections of *P. gingivalis* resulted in translocation of bacteria to the brain, increased Aβ1-42 secretion and AD-like neurodegeneration^36,37^. Interplay between Aβ and *P. gingivalis* also occurs outside the central nervous system. Notably, Aβ40/42 was detected in the gingival crevicular fluid of AD patients with periodontitis, where it aggregated within the gingival biofilm^38,39^.

Although *P. gingivalis* is considered a pathogen of interest in AD, the impact of its biofilm on Aβ has yet to be characterized. This multicellular structure is important, since bacterial biofilms were recovered as part of Aβ plaques^40–42^. A better understanding of the interaction between these two components will provide significant insights into how *P. gingivalis* could be a risk factor for AD and how it can persist despite Aβ’s antimicrobial activity. In this study, we co-cultivated *P. gingivalis* under biofilm-inducing conditions alongside Aβ1-40 and Aβ1-42 (Aβ40/42) and observed that the interaction between Aβ and the biofilm of *P. gingivalis* depends on the peptide subtype.

## Results

### Aβ influences P. gingivalis biofilm formation

Previous reports have identified Aβ40/42 as an antimicrobial peptide against multiple bacterial species such as *Escherichia coli*, *Enterococcus faecalis*, and *Staphylococcus epidermidis*^27^. To determine if Aβ40/42 would impact *P. gingivalis* strain 33277 viability or biofilm formation, *P. gingivalis* was co-incubated for 24 h with various concentrations of Aβ40/42 under biofilm-inducing conditions. These Aβ concentrations were chosen to reflect a microenvironment in which bacteria are not abundant but Aβ40/42 is, such as in the inflamed part of the brain, and they were also used previously^27,31^. After the co-incubation, we quantified biofilm formation using a crystal violet assay. Surprisingly, the two Aβ peptides tested had different impacts on biofilm formation. High concentrations of Aβ1-40 lowered biofilm production at 25 (55% decrease) and 12.5 μg/mL (34% decrease) (Figure 1). Meanwhile, incubation with Aβ1-42 had the opposite effect and resulted in a dose-dependent, significant increase in biofilm formation, peaking between 12.5 (56% increase) and 25 μg/mL (46% increase) (Figure 1). In parallel, we examined whether Aβ40/42 impacted *P. gingivalis* viability in planktonic cultures using LIVE/DEAD™ assays followed by flow cytometry analysis. Interestingly, the highest concentration of Aβ40 or Aβ42 did not affect *P. gingivalis* (Figure S1).

**Figure 1.**
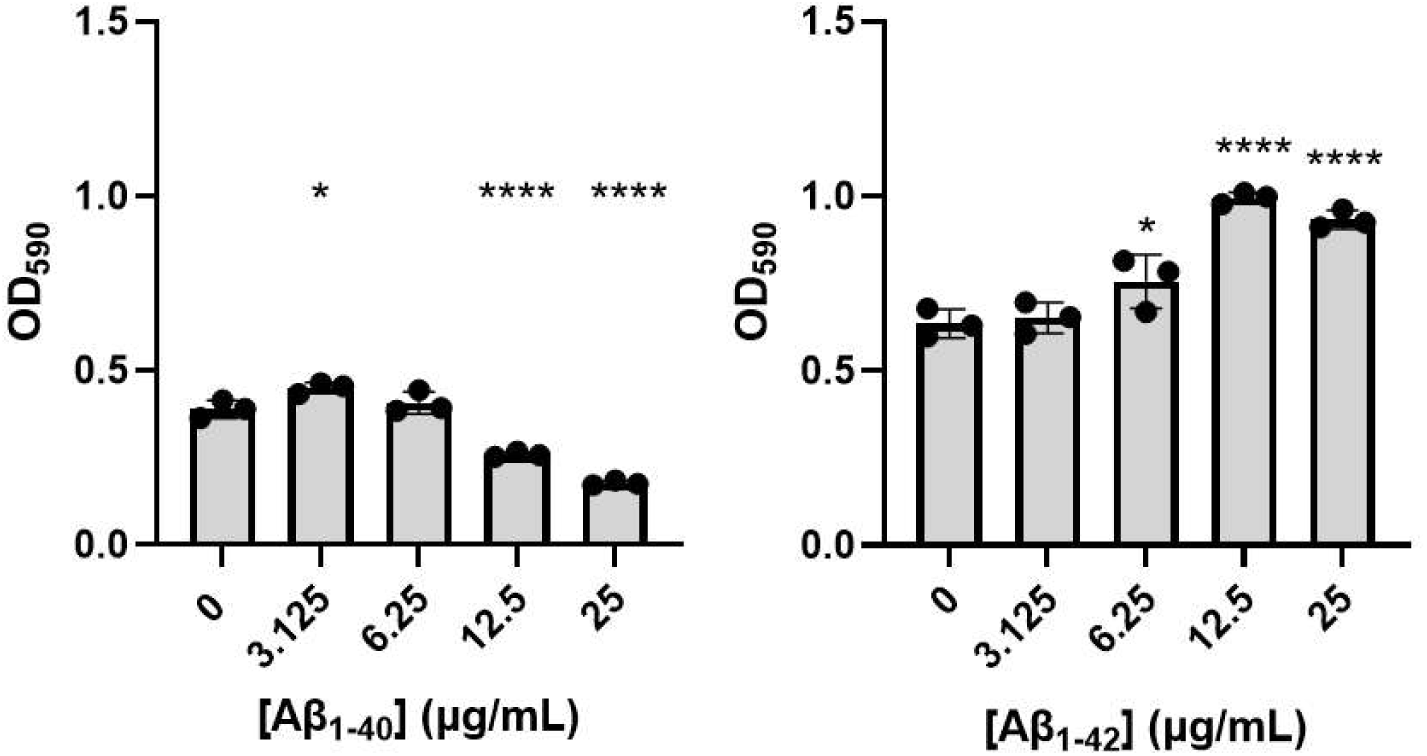
Influence of AB40;42 on *P gingivalis* biofilm growth. Biofilm production of *P gingivalis* in the presence of varying concentrations of AP1-4o (Left) or AB1-42 (Right) in µg/mL, as measured by crystal violet staining after 24 h. All statistical analysis was performed using a Brown-Forsythe andWelch’ss analysis of covariance (ANOVA), * (P:S0.05), **** (P:S0.0001) indicate a significant difference from the control (0 µg/mL). Error bars represent the standard deviation (SD).

Since crystal violet is a non-specific dye, the impact of Aβ40/42 on biofilm formation could be due to a change in the amount of extracellular matrix or bacteria inside the biofilm. To discriminate between these two possibilities, we used differential fluorescence staining to quantify the abundance of each component. We used the DNA-binding fluorescent molecule 4’,6-diamidino-2- phenylindol (DAPI) to mark the cells. Thioflavin-T (ThT) was used as a general protein marker, as a proxy for proteins within the extracellular matrix^43^. Despite being quite specific to amyloid fibers, ThT was shown to be fluorescent in the *P. gingivalis* biofilm and in cell-free biofilm fragments (Figure S2). Finally, the matrix exopolysaccharides were marked using a fluorophore-bound lectin ConA^44^. These markers were used to stain *P. gingivalis* biofilms cultivated in the presence of 25 μg/mL Aβ1-40 or Aβ1-42, and fluorescence was quantified to determine if the abundance of cells, proteins or exopolysaccharides varied during biofilm formation. For Aβ1-40, we observed a significant decrease in the fluorescence intensity of each marker compared to the control with no Aβ (Figure 2A). Indeed, direct observation of the biofilm by confocal imaging (Figure 2B) confirmed that in the presence of Aβ1-40, the biofilm of *P. gingivalis* was very sparse and organized in small aggregates.

**Figure 2.**
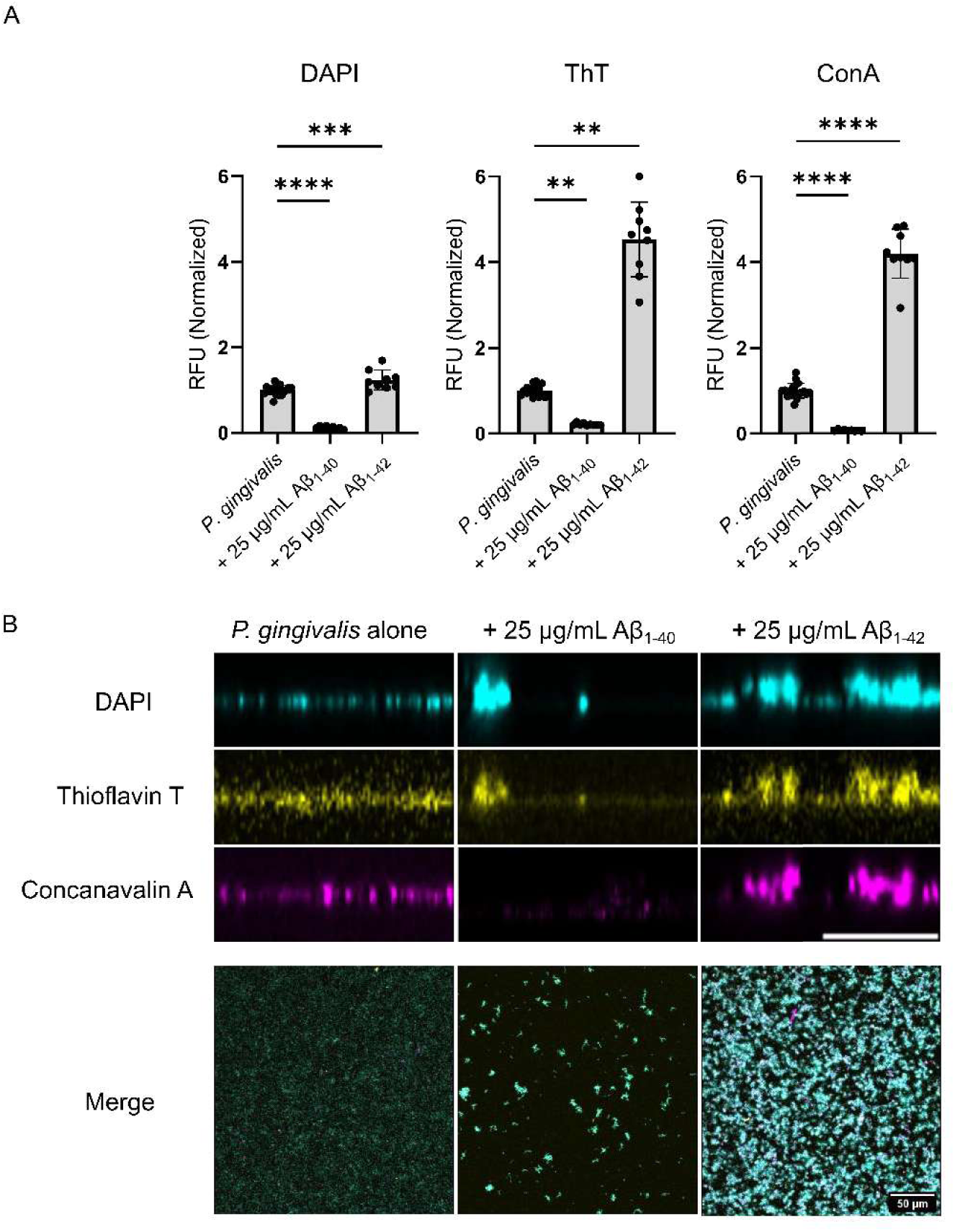
Influence of AP1-4o and AP1-42 on *P gingivalis* biofilm composition and thickness. (A) Fluorescence intensity fold change ofDAPI, Thioflavin T and Concanavalin A staining on *P gingivalis* biofilm cultivated with 25 µg/mL AP40;42 for 24 h. Values for the control were pooled from the individual assays on AP1-4o and AP1-42 and normalized against the mean. Statistical analysis was performed using ANOVA for DAPI and ConA and Kruskall-Wallis for ThT, ** (P:S0.005), *** (P:S0.001), **** (P:S0.0001). Error bars represent SD. (B) Cross-sections of *P gingivalis* biofilm cultivated with 25 µg/mL AP40;42 for 24 h and visualized with DAPI, ThT and ConA. Scale Bar= 10 µm. Accompanied by the merge of the three channels on a front-view. Scale Bar = 50 µm.

Meanwhile, biofilms of *P. gingivalis* formed with Aβ1-42 exhibited a notable increase for each marker compared to the control, the degree of which varied among markers. While the increase in intensity associated with DAPI (∼1.25 fold) was significant, it was much lower than the fluorescence increases associated with ThT (∼4.5 fold) and ConA (∼4.2 fold). This result suggests that the effect of Aβ1-42 on the biofilm was due to a strong stimulation of the production and secretion of the extracellular matrix (Figure 2A). Of note, the sharp increase in the ThT fluorescence intensity in the presence of Aβ1-42, which surpasses the increase observed with ConA, could be partly due to ThT binding to the aggregated Aβ within the biofilm. Confocal imaging of biofilm cross-sections showed a thicker community when the biofilm was formed in the presence of Aβ1-42 (Figure 2B), resulting in a higher signal in all channels and a visibly increased biomass.

### Aβ aggregates within the P. gingivalis biofilm

Previous studies identified biofilm components in the brain of AD patients, interwoven with Aβ plaques^40,41^. We hypothesized that, similarly, Aβ aggregates could be embedded in the biofilm of *P. gingivalis*. Therefore, we imaged biofilms of *P. gingivalis* formed in the presence of 5 μg/mL of Aβ40/42. This concentration did not strongly disrupt biofilm formation, similarly to 3.125 and 6.25 μg/mL (see Figure S3), which was important since our goal was to visualize if Aβ40/42 could be embedded in the biofilm. Matrix components were stained as described in Figure 2, and Aβ40/42 was detected using the MOAB-2/6C3 monoclonal antibody coupled to the Alexa-555 fluorophore. Interestingly, confocal imaging revealed that aggregates of Aβ were present as inclusions in the extracellular matrix of *P. gingivalis* after 24 hours. Aβ1-42 within the *P. gingivalis* biofilm was more abundant than Aβ1-40, which was present in only a few instances (Figure 3A). Quantification of the Alexa-555 in the biofilm confirmed that the accumulation of Aβ1-42 was significantly higher than Aβ1-40 (Figure 3B). We then assessed if Aβ1-42 aggregation was more associated with cells (DAPI) or the extracellular matrix (ConA). Mander’s co-localization correlation analysis showed that the signal for Aβ1-42 co-localized significantly more with the Concanavalin-A channel than DAPI (Figure 3C).

**Figure 3.**
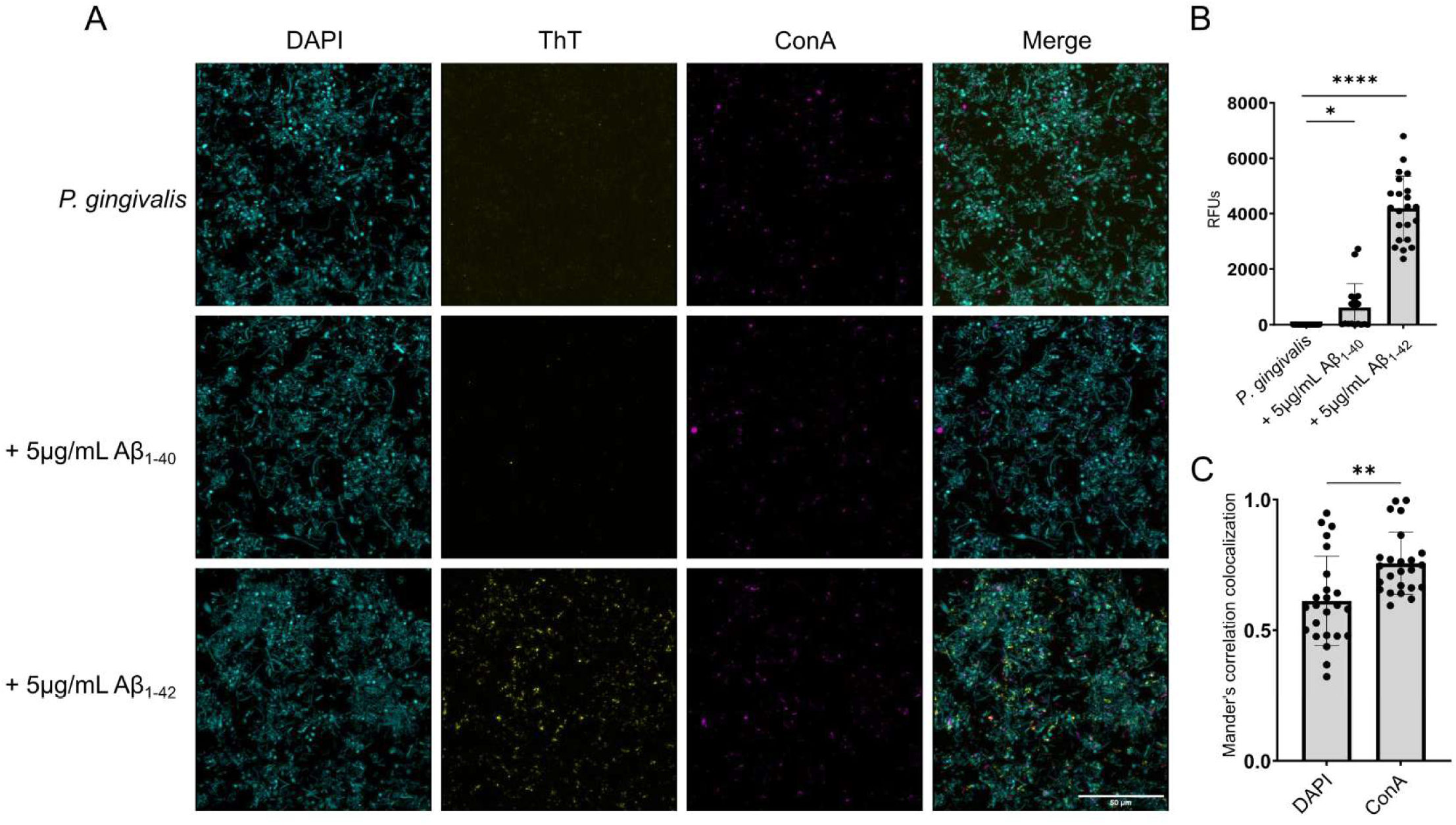
Aggregation of AP40/42 within the biofilm of *P. gingivalis.* (A) Confocal imaging ofAPi. 40 or APi.42 (Yellow; ThT) aggregated within the biofilm of *P. gingivalis* after 24 h. Cells are in blue (DAPI) and biofilm is in magenta (ConA). (B) Quantification of the fluorescence of AP1-40 or APi- 42 aggregated within the biofilm of *P. gingivalis.* Statistical analysis was performed using ANOVA, *(*P* ≤ 0.05), ****(*P* ≤0.0001) (C) Manders colocalization correlation of Ap with DAPI (Cells) and ConA (Biofilm). Student’s T Test. **(*P* ≤0.005)

### P. gingivalis biofilm influences Aβ aggregation

Since Aβ aggregates were visible within the biofilm of *P. gingivalis,* we hypothesized that some biofilm components could interact with Aβ peptide and influence its aggregation rate. We first separated the extracellular biofilm matrix from the bacterial cells to produce nanometric fragments of biofilm (hereafter named seeds). These seeds were then incubated with monomeric Aβ40/42 in a 1:20 ratio in the presence of ThT, which binds to β-fold structures formed during amyloid aggregation. This binding leads to an increased fluorescence from ThT, allowing the observation of Aβ aggregation in real time. Real-time fluorescence curves were then analyzed by assessing the aggregation constant (k), the aggregation half-time and the length of the lag phase. For Aβ1-40, we observed a significant decrease in lag phase when the peptide was incubated with seeds, which could also be observed when looking at the resulting aggregation curve (Figure 4A). We also observed a non-significant, but still sizeable, decrease in half-time. Interestingly, incubation of monomeric Aβ1-40 with seeds also induced a 3-fold reduction in the aggregation constant of the peptide. These results indicate that although the presence of seeds reduced the time necessary for fibrillation of Aβ1-40, the actual aggregation rate was slower. For Aβ1-42, incubation with seeds had no impact on its aggregation onset or rate (Figure 4B).

**Figure 4.**
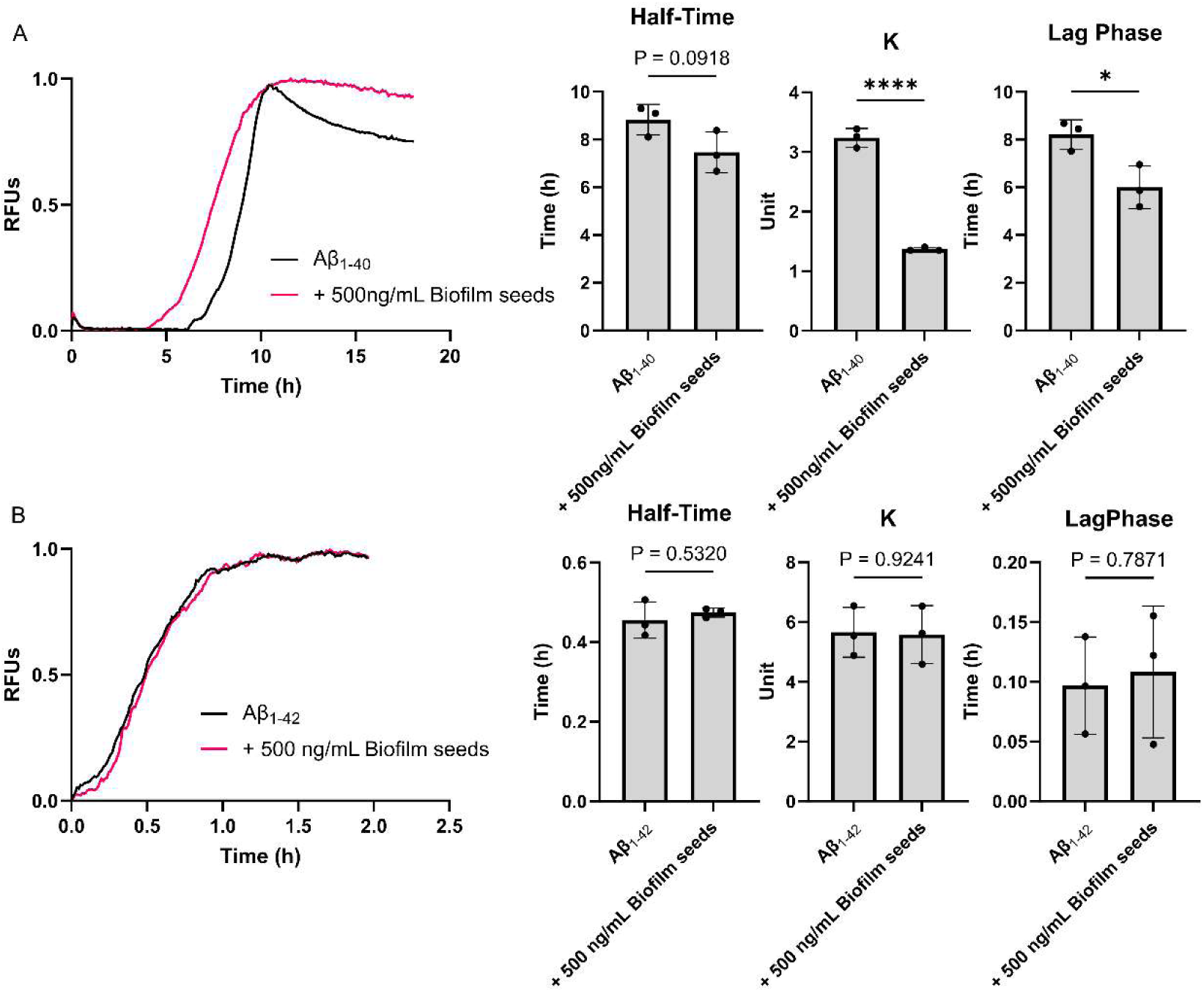
Aggregation kinetic of monomeric APmo (A) and AP1-42 (B) incubated with biofilm seeds from *P. gingivalis.* Curves are the average of 3 technical replicates. Half time (Left), Aggregation constant (K, Middle) and Lag Phase (Right) were extracted from individual curves. Statistical analysis was performed with Student’s T test, *\*(P<0.05),* ****(P<0.0001). Error bars represent SD.

We then used Atomic Force Microscopy (AFM) to assess if the increase in Aβ1-40 aggregation in the presence of seeds resulted in a change in the overall structure of fibrils. As shown in Figure 5, Aβ1-40 alone organized in lengthy and complex networks of interconnected fibrils (Figure 5A). However, in the presence of seeds, Aβ1-40 formed tight and isolated bundles of thick fibrils. This seemingly higher aggregative state might explain the decrease in lag phase necessary for Aβ aggregation observed in Figure 4A. For Aβ1-42, the aggregation pattern between the peptide alone and with seeds exhibited no significant difference, which correlates with results obtained in ThT assays (Figure 5B).

**Figure 5.**
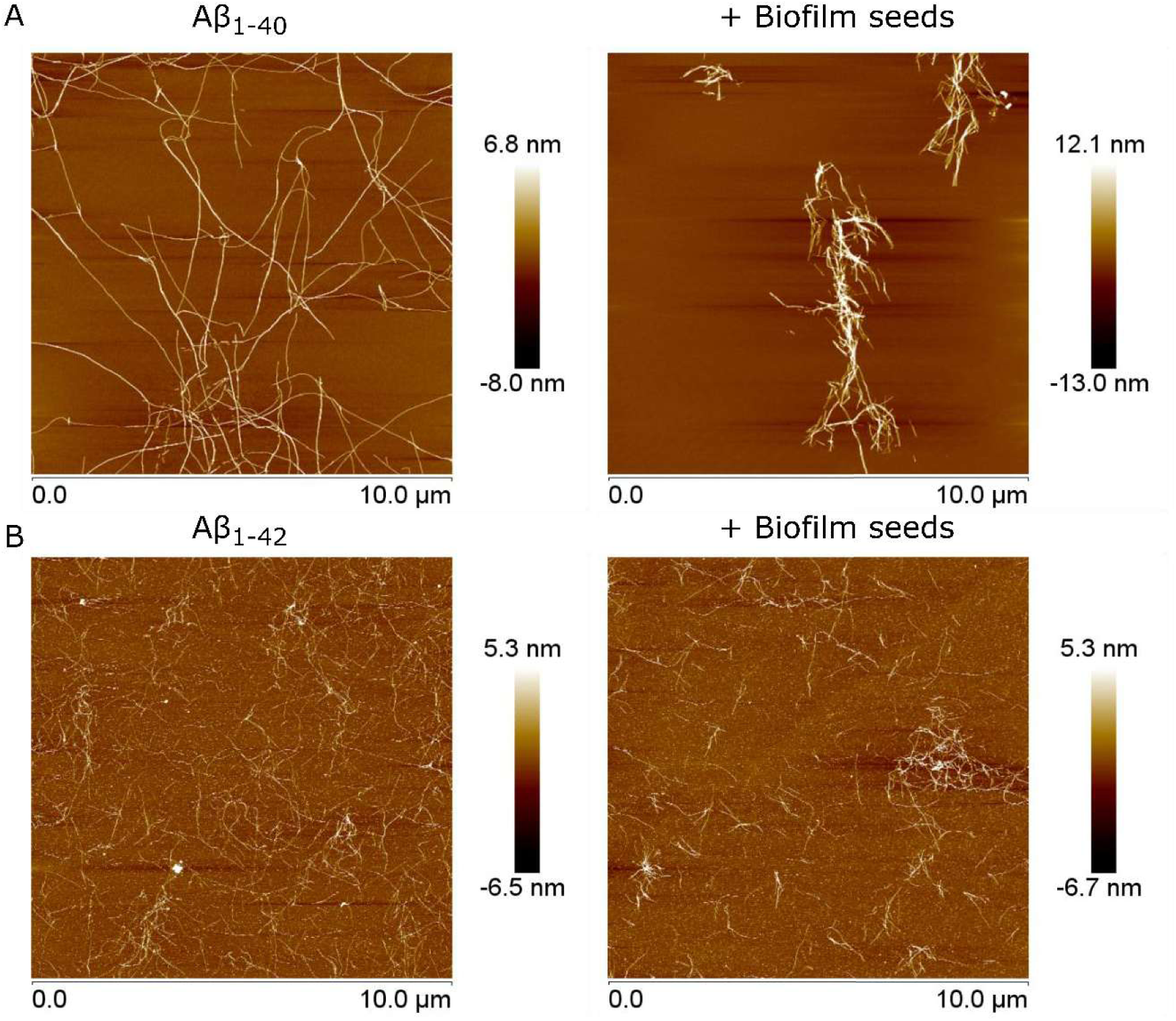
Aggregates from AP1-40 (A) or Api-42 (B) incubated alone or with biofilm seeds from *P. gingivalis.* Visualized through AFM using height sensors. Images are representative of 12 technical replicates over 3 biological replicates.

### Gingipains can break down Aβ

*P. gingivalis* produces gingipains, large-spectrum proteases able to break down antimicrobial peptides^45^. Since the biofilm of *P. gingivalis* was resistant to Aβ1-42 antibacterial properties, gingipains, which are abundant in the biofilm, could be involved in limiting its toxicity. We therefore assessed the proteolytic activity of gingipains on Aβ40/42. We first produced gingipain- rich stationary phase supernatants from a 48-hour culture of *P. gingivalis* 33277, of the deletion mutants Δ*rgp*AΔ*rgpB* (KDP112), Δ*kgp* (KDP129), and of the triple mutant Δ*rgpAΔrgpB*Δ*kgp* (KDP128). All supernatants were incubated with 5 μg/mL Aβ40/42 for 16 hours and then spotted on PVDF membranes for dot blot assays. As shown in Figure 6A, both Aβ1-40 and Aβ1-42 were degraded when incubated with supernatants from *P. gingivalis* 33277, demonstrating that the bacterial supernatant can break down the peptide. Exposure to the supernatant of Δ*rgpAΔrgpB*Δ*kgp* did not affect the peptide, suggesting that gingipains are responsible for Aβ40/42 degradation. Interestingly, the supernatant produced by the Δ*rgpA*Δ*rgpB* strain also did not break down Aβ40/42 whereas the supernatant from Δ*kgp* did (Figure 6A). Interestingly, gingipains could only digest monomeric Aβ40/42, while fully aggregated fibrils were unaffected (Figure 6B). Aβ fibrils are notoriously complex to break down, requiring a multi-step process that includes various proteases^46^. In our case, gingipains alone were insufficient to degrade pre-assembled Aβ complexes fully.

**Figure 6.**
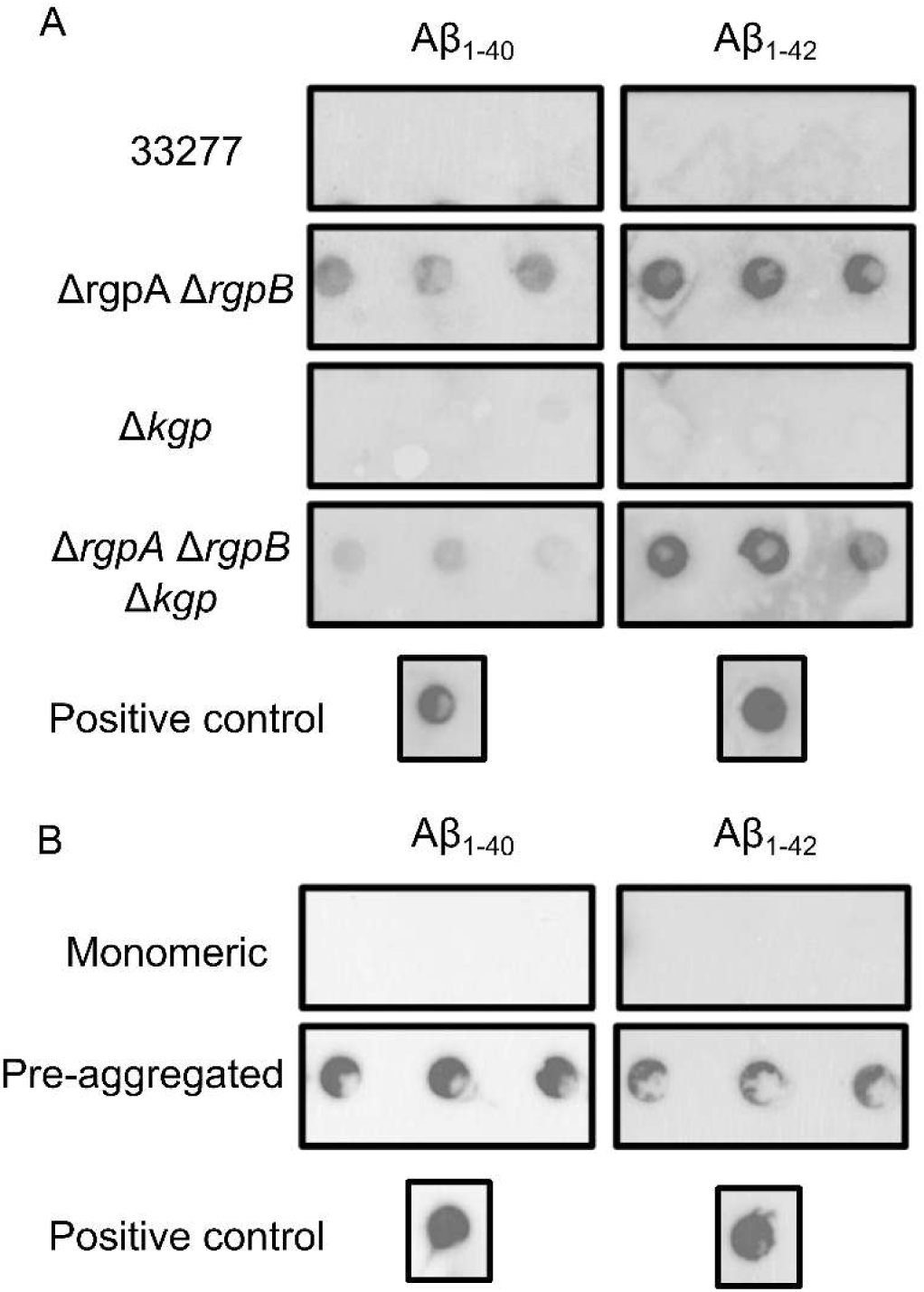
Degradation of AP40/42 by gingipains from *P. gingivalis.* (A) Monomeric APmo or APm2 incubated with supernatant from *P. gingivalis* WT ATCC 33277, *Δkgp. ΔrgpA ΔrgpB.* and *ΔrgpA ΔrgpB Δkgp.* (B) Monomeric and pre-aggregated (24 h at 37 °C) of APmo or APm2 incubated with supernatant from *P. gingivalis* ATCC 33277. Results are representative of three biological replicates. Positive controls represent Ap without supernatant from *P. gingivalis.*

## Discussion

While the link between AD and *P. gingivalis* was investigated over the past 10 years, no study has examined the specific impact of the biofilm on the AD hallmark that is Aβ^19,35,37,47–53^. Using a simplified model, we were able to observe a bidirectional relationship that was unique to each peptide. Indeed, both Aβ impacted *P. gingivalis* biofilm production differently, but the biofilm only influenced Aβ1-40 aggregation.

We first determined that Aβ1-40 and Aβ1-42 had different effects on *P. gingivalis*. Biofilm production was decreased when exposed to increasing concentrations of Aβ1-40, while Aβ1-42 increased the abundance of the extracellular matrix. These results are surprising considering that Aβ1-42 is more toxic than Aβ ^54^. Importantly, Aβ did not display antimicrobial activity on *P. gingivalis*, since it stimulated biofilm formation and did not impact the viability of planktonic cells. This lack of antagonism might be due to the abundance of proteinases secreted by *P. gingivalis*, as discussed later. Considering that Aβ1-42 stimulated biofilm formation in a dose-dependent manner (Figure 1) and that there is a lack of interaction between Aβ1-42 and the biofilm seeds (Figure 4), the increase in extracellular matrix could be the result of a high quantity of Aβ aggregates embedded within the biofilm (Figure 3).

In contrast to Aβ1-42, high concentrations of Aβ1-40 resulted in decreased biofilm formation. These observations are not unlike observations made on *P. aeruginosa,* for which the presence of Aβ1-42 decreased biofilm production^55,56^. Interestingly, this bacterium relies on the presence of the functional amyloid protein FapC as part of its extracellular matrix^57,58^. FapC can cross-seed Aβ1-42 and form unproductive heterogeneous polymers, leading to a disruption of the biofilm^59^. While there are currently no known functional amyloid proteins in *P. gingivalis* biofilm, ThT assay (Figure 4) and AFM imaging (Figure 5) showed that Aβ1-40 aggregation pattern differed in the presence of biofilm seeds. These results suggest the existence of a co-aggregation phenomenon between Aβ1-40 and a component of the biofilm matrix. This co-aggregation appears unproductive, leaving clumps of matrix rather than a fully formed biofilm (Figure 2), similar to what was observed with *P. aeruginosa*^55^.

In contrast, ThT assays and AFM imaging showed no interaction between Aβ1-42 and *P. gingivalis* biofilm. This result could stem from a faster Aβ1-42 self-assembly kinetic compared to its potential interaction with a biofilm component. Indeed, Aβ1-42 possesses a high aggregation potential and might favor homogenous aggregation instead of cross-interactions (reviewed in^60^), thus not interfering with *P. gingivalis* biofilm matrix polymerization (Figure 2).

Aβ40/42 inclusions within the biofilm appeared unrelated to co-aggregation capacity, since we observed both Aβ1-42 and Aβ1-40 aggregates embedded within the matrix. These inclusions seemed to result from Aβ40/42 aggregates trapped in the extracellular matrix following deposition^61^, which would also explain the co-localization with exopolysaccharides. Strikingly, only a small number of Aβ1-40 inclusions were present compared to Aβ1-42. The outcome might be due to the slower aggregation rate of Aβ1-40 and the presence of gingipains produced by *P. gingivalis*. Indeed, gingipains can degrade monomeric, but not pre-aggregated Aβ (Figure 6), since the hydrophobic structures between β-sheets of amyloid oligomers make them resistant to enzymatic breakdown^62^. Thus, we hypothesize that there are fewer Aβ1-40 inclusions embedded in the biofilm because Aβ1- 40 exists as monomers and low-weight oligomers for a longer duration than Aβ1-42, providing more opportunity for degradation by gingipains.

Proteolysis of antimicrobial peptides, including human peptides, is a key strategy of *P. gingivalis* to evade the host’s immune response^63,64^. Interestingly, we observed that deletion of arginine-specific gingipains (RgpA/RgpB) but not of the lysine-specific gingipain (Kgp) negated Aβ degradation, despite lysine residues being present in both subtypes of Aβ. These two types of gingipains are known to have differences in activity and inhibitory conditions, which might explain why only RgpA/RgpB appeared to degrade Aβ40/42 in our conditions ^66,67^. Further experiments are required to determine if it is also the case *in situ*.

Aβ1-42 differs from Aβ1-40 only by the addition of an isoleucine and an alanine in C-terminal, which increase Aβ1-42 overall aggregation rate into protofibrils and fibrils^68,69^ compared to Aβ1-40. These properties are likely sufficient to explain the difference observed in our study. Indeed, the high aggregation rate of Aβ1-42 and its stability in bigger oligomers would limit its interaction with *P. gingivalis* biofilm components and its degradation by gingipains. Proteolysis-resistant oligomers and fibrils would then integrate the biofilm and increase its overall biomass. In opposition, Aβ1-40 exists in an equilibrium of monomers and low complexity oligomers for a substantial time before assembling into fibers, allowing for more interactions with biofilm matrix components. These interactions would be unproductive for both partners, Aβ1-40 fibrils displaying unusual bundles, and the biofilm matrix being unable to fully polymerize.

*P. gingivalis* biofilm components were identified in Aβ plaques isolated from AD patients, suggesting that a co-aggregation phenomenon with Aβ1-40 and/or an integration of Aβ1-42 aggregates in the matrix could occur within the brain^40,41^. Indeed, modulation of Aβ aggregation by physical interaction with viral or bacterial proteins is one of the core elements that could explain how microorganisms would contribute to AD development^55,70,71^. *P. gingivalis* biofilms are also routinely observed on teeth^72^, and consequently, our observations demonstrate a potential interaction between *P. gingivalis* and Aβ outside of the central nervous system (CNS). Aggregates of Aβ were observed on the surface of teeth and embedded within the oral biofilm of patients with AD^39^, which concurs with our observations. These non-CNS interactions are important, since peripheral infections of *P. gingivalis* were also linked to AD-like dementia in mice^37,48^.

The *P. gingivalis* extracellular matrix component that interacts with Aβ1-40 and alters its aggregation dynamics has yet to be identified. Additionally, FimA subtypes and capsular polysaccharides and hemagglutinin expression vary between strains, which may influence the impact of biofilms on Aβ ^73,74^. Nonetheless, by shedding light on the interaction between Aβ and *P. gingivalis*, our study offers a new perspective on this bacterium’s potential role in the onset of AD. While the exact etiology remains unclear, a deeper understanding of the mechanisms underlying key risk factors such as periodontitis can provide valuable knowledge for prevention and treatment strategies.

## Methods

### Aβ preparation

Aβ1-40 and Aβ1-42 (Genscript, NJ, USA) were prepared as previously described^75^. Briefly, 1 mg of lyophilized Aβ40/42 was dissolved in a solution of NH4OH 10% (w/v) to 0.5 mg/ml. The peptides were incubated 10 minutes at room temperature before being sonicated for 5 minutes and aliquoted in microcentrifuge tubes. The aliquots were then lyophilized overnight and stored at -80 ᵒC. While lyophilization should get rid of the NH4OH, we also verified that it did not influence biofilm formation at the maximum working concentration (Figure S4).

### Strain and media

*Porphyromonas gingivalis* ATCC 33277, KDP112, KDP128 and KDP129 were generously provided by Daniel Grenier (Université Laval, Québec). Strains KDP112 (Δ*rgpA ΔrgpB*), KDP 129 (*Δkgp*) and KDP128 (*ΔrgpA ΔrgpB Δkgp*) are gingipain deletion mutants in the *P. gingivalis* ATCC 33277 genetic background^76,77^. Cultures were routinely maintained in Tryptic-Soy broth supplemented with hemin and vitamin K (TSBHK; 17 g/L casein peptone, 2.5 g/L K2HPO4, 2.5 g/L glucose, 5 g/L NaCl, 3 g/L soy peptone, 5 μg/mL hemin and 1 μg/mL vitamin K) in a Type-B vinyl COY anaerobic chamber with the following gas mix: 80% N2:10% CO2:10% H2, in static conditions. For biofilm-inducing conditions, *P. gingivalis* was cultured statically in TSBHK supplemented with Tryptone 1% (w/v) (TSBHKT)^78^.

### Crystal violet assay

Crystal violet assays were used to assess biofilm production under different concentrations of Aβ ^79^. First, 36 h cultures of *P. gingivalis* were centrifuged at 3200 xg for 20 minutes and resuspended in TSBHKT. Aβ aliquots were resuspended in 60 mM NaOH and diluted to the proper concentration in the same medium. Biofilm cultures were then performed in polystyrene 48-well plates containing 200 mL of TSBHKT inoculated with *P. gingivalis* at OD600= 0.005 and incubated for 24 h at 37 ᵒC. Of note, this concentration of *P. gingivalis* was selected after optimisation for the most robust biofilm formation in our conditions. The remaining biofilm was washed with phosphate-buffered saline (PBS; 0.2 g/L KCl, 1.42 g/L Na2HPO4, and 0.24 g/L KH2PO4, 0.8 g/L NaCl, pH 7.4) before being covered with 200 μL of 0.01% w/v crystal violet for 20 minutes at room temperature. Each well was then washed with sterile deionized water to remove excess dye and resuspended acetic acid 33% (v/v) before being quantified at OD590.

### Imaging of biofilm components

For confocal imaging of biofilm with Aβ40/42, *P. gingivalis* was inoculated at OD600= 0.005 in 200 μL TSBHKT + 25 μg/mL Aβ in flat clear-bottom microscopy 96-well plates (Ibidi, Gräfelfing, Germany) for 24 h at 37 ᵒC. Supernatants from the cultures were discarded and the remaining biofilms were washed with PBS. The biofilms were then fixed for 7 minutes in 200 μL of 4% (v/v) paraformaldehyde and then blocked with 200 μL of PBS + 2% (w/v) bovine serum albumin for 1 h at room temperature. For the staining of biofilm components under high Aβ concentrations (Figure 2), we used Concanavalin A-Alexa647™ (ConA) (1 h at room temperature, 50 μM; Thermo Fisher, USA), Thioflavin T (ThT) (30 minutes at room temperature, 100 μM; Millipore Sigma, USA) and 4′,6-diamidino-2-phenylindole (DAPI) (30 minutes at room temperature, 1 μg/mL; Millipore Sigma, MA, USA). Pictures of the biofilms were taken using an Olympus FV3000 (Olympus, JP) confocal microscope at 20x magnification (PlanApo objective [20X/0.75 NA (Numerical aperture)]) with the following laser settings: 405 nm (DAPI), 445 nm (ThT) and 640 nm (ConA). Laser intensity and sensor sensitivity were the same for all conditions. Image analysis used at least 25 images per condition, spread over 3 biological replicates. To observe Aβ aggregation within the biofilm (Figure 3), we used Concanavalin A-Alexa647™ (1 h at room temperature, 50 μM, Thermo Fisher, USA), 6C3 anti-amyloid beta peptide monoclonal antibody (1 h at room temperature, 1:1000, Millipore Sigma, MO, USA), Goat anti-Mouse IgG (H+L) Cross-Adsorbed Secondary Antibody, Alexa Fluor™ 555 (1 h at room temperature, 2 μg/mL, Thermo Fisher, USA) and DAPI (30 minutes at room temperature, 1 μg/mL, ThermoFisher, USA). The images were produced with the same confocal microscope at 40x magnification (PlanApo objective [40X/0.95 NA]) with the following laser settings: 405 nm (DAPI), 561 nm (AlexaFluor™ 555) and 640 nm (ConA). Laser intensity and sensor sensitivity were the same for all conditions.

### Microscopy analysis

For the fluorescence intensity measurements, Z-stack confocal images were initially preprocessed using Fiji. For each image, Z sum intensity projection was applied, followed by a rolling ball background subtraction. Subsequently, whole image intensity measurements were obtained using CellProfiler image analysis software. For the colocalization analysis, Fiji was used to perform maximum intensity projection on z-stacks images. Pixel colocalization was then measured on projection images using the Measure Colocalization module of CellProfiler 4.2.6^80^. Manders coefficient was used to measure correlation between different fluorescence channels^81^.

### Biofilm seeds preparation

Biofilm fragment preparation was done as described elsewhere^70^. Briefly, multiple wells in a 96- well plate were inoculated with *P. gingivalis* 33277 in 200 μL of TSBHKT at OD600=0.005. The biofilm was allowed to form in static conditions at 37 ᵒC for 24 h. The biofilm was then scooped in sterile PBS and vortexed at low intensity for 10 minutes with 25 μm glass beads before being spun down at 3200 xg for 10 minutes. The supernatant was then collected and quantified using Pierce™ BCA Protein Assay Kit (Thermo Scientific, MA, USA). Of note, proteomic analysis of these biofilm seeds confirmed the presence of all major and minor fimbria subunits, and CFU quantification reveals 10^3^ less cells in seeds than in biofilm (Figure S5).

### ThT assays

For ThT assays, Aβ aliquots were resuspended in 10 μL of 60 mM NaOH and incubated 5 minutes at room temperature. They were then diluted to 10 μg/mL in PBS containing 30 μM ThT and distributed in 200 μL in black 96-well polystyrene plaques (Corning, NY, USA) in presence of 500 ng/mL biofilm seeds. The assay wells were surrounded by two layers of wells containing water to account for evaporation. Plaques were read every 5 minutes over a 20-hour period using a TECAN SPARK plate reader at an excitation wavelength of 440 ± 20 nm and an emission wavelength of 485 ± 20 nm under constant agitation at 37 ᵒC. Resulting curves were analyzed to assess kinetic values as previously published elsewhere^82^. Briefly, ThT results were fitted to a sigmoidal curve (Eq.1) where k is the apparent aggregation constant, t1/2 is the time to reach half the fluorescence.

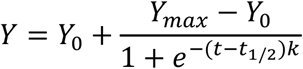

From this equation, it is also possible to measure the lag phase using Eq.2.

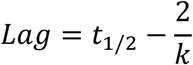

### Atomic force microscopy (AFM)

Experiments leading to AFM were performed in the same way as ThT assays. Briefly 100 ug/mL solutions of Aβ40/42 were incubated in PB buffer (0.2 g/L KCl, 1.42 g/L Na2HPO4, and 0.24 g/L KH2PO4, pH 7.4) with 0.5 ug/mL of biofilm fragments. Aβ40/42 samples were incubated in static conditions at 37 ᵒC for 3 hours for Aβ1-40 and 1 hour for Aβ1-42 before being adsorbed in a humid atmosphere on a clean mica surface (Electron Microscopy Sciences, PA, USA) for 30 minutes. Following adsorption, mica slides were washed twice in ddH2O before being dried under a nitrogen stream. AFM was performed using a Veeco Dimension Icon™ microscope (Bruker, USA) using ScanAsyst air probes (Bruker, resonance frequency 70 KHz, spring constant 0.4 N/m, tip nominal radius 2 nm). For each condition, 15 images were captured over 3 biological replicates for an area size of 10 μm/10 μm.

### Gingipain assays

*P. gingivalis* WT and the gingipains mutants were statically grown in TSBHK for 48 h and adjusted to OD600=1. Cells were then removed by centrifugation at 3200 xg for 20 minutes. 10 μL of *P. gingivalis* supernatant was then mixed with 190 μL of a 5 μg/mL Aβ40/42 solution in PBS. Samples were incubated at 37 ᵒC for 16 h before being analyzed by dot blotting. 20 μL of sample was spotted on a PVDF membrane and immunoblotted using MOAB-2 6C3 anti-amyloid beta peptide primary antibody (1 h at room temperature, 1:2500, Sigma, MO, USA) and Peroxidase AffiniPure Goat Anti-Mouse IgG (H+L) (1 h at room temperature, 1:5000; Jackson ImmunoResearch, PA, USA). Membranes were revealed using Clarity Max ECL Western Blotting Substrates (Biorad, USA) and imaged through Biorad ChemiDoc using chemiluminescence. To observe the difference between non-aggregated and aggregated Aβ40/42, a sample of each Aβ subtypes was initially incubated in PBS at 37°C for 24 hours. This sample was compared to a fresh aliquot of Aβ and analyzed using the method described above.

### Statistical analysis

Statistical analyses were performed in Graph Pad Prism 10. Comparisons were done using Student’s t-test, one-way analysis of variance (ANOVA) or Kruskal-Wallis tests, all with 95% confidence intervals. Normality was assessed using Shapiro-Wilke’s normality test and each result has been replicated in at least three independent biological replicates.

## Acknowledgement

We would like to thank Dr Grenier for the kind gift of strains, J. Beaudin for the critical review of the manuscript, and members of the Beauregard lab for insightful discussions. This project was financed by FRQ AUDACE grant 2019-AUDC-263669, the CR-CHUS support funds, the Ministère des Relations internationales et de la Francophonie du Québec and by a PhD fellowship (FRQS) to D. Dumoulin.

